# Bacterial supplementation shapes honey bee gut microbiota, metabolism and proteome under controlled and field conditions

**DOI:** 10.1101/2025.11.20.689477

**Authors:** Saetbyeol Lee, Pavel Dobes, Anna Mascellani Bergo, Jacek Marciniak, Jana Hurychova, Sara Sreibr, Martin Kamler, Vojtech Purnoch, Jiri Killer, Lucie Hlinakova, Dalibor Titera, Ondrej Cinek, James C. Carolan, Pavel Hyrsl, Jaroslav Havlik

## Abstract

Gut bacteria are essential to honey bee (*Apis mellifera*) health, supporting digestion, immunity, and resilience against stressors. Probiotic-based strategies have been proposed to enhance core gut symbionts, yet the underlying mechanisms and influences of environmental context on these effects are not fully understood.

This study examined the influence of gut-derived bacterial supplementation on the honey bee gut microbiome, proteome, and metabolome across three conditions: (i) controlled cages, (ii) semi-controlled cages allowing contact with nestmates, and (iii) field conditions. Treatment groups received a bacterial supplement containing *Lactobacillus helsingborgensis, Lactobacillus apis, Bifidobacterium choladohabitans*, and *Bifidobacterium polysaccharolyticum,* while control groups received only a sucrose solution. Gut samples from 10-day-old bees were analyzed.

Supplemented bees showed strong gut colonization by *Bifidobacterium* and *Lactobacillus,* especially under controlled and semi-controlled conditions. In the field, *L. helsingborgensis* remained significantly enriched in the treatment group, indicating ecological resilience. Proteomic changes in supplemented bees included an increased abundance of major royal jelly protein precursors and mitochondrial proteins, alongside the decreased abundance of ribosomal and translation-related proteins involved in peptide biosynthesis and cellular protein quality control. Metabolomic analysis revealed reproducible shifts across all three conditions. Treatment groups showed higher concentrations of microbial fermentation products (acetate, succinate) and neuroactive compounds (ornithine, γ-aminobutyrate), while sucrose, *N*-acetylglucosamine, and uridine that were constantly lower compared to controls. No significant changes were observed in body or gut weight.

These findings highlight reproducible, context-dependent effects of bacterial supplementation on honey bee gut physiology and provide mechanistic insights for microbiome-based interventions in pollinator health.

**Importance:** Honey bees are essential pollinators whose health is influenced by their gut microbiome. Probiotic applications aimed at improving gut health have been proposed, yet outcomes remain inconsistent and vary across settings. Results from laboratory experiments often differ from those observed under field conditions, making it difficult to understand the complex dynamics of eusocial insect colonies. Here, we evaluate honey bee gut-derived bacterial supplementation across controlled, semi-controlled, and field settings using bacterial profiling, proteomic, and metabolomic analyses. We demonstrate that bacterial supplemented groups consistently reshape gut community composition and modulate host physiological processes, but in a context-dependent manner. These results provide a unified understanding of how microbial interventions function at colony and individual levels, guiding the rational design of probiotic strategies to support honey bee health under realistic conditions.

## 1. Introduction

Honey bee (*Apis mellifera*) colonies are experiencing widespread declines, driven by multiple interacting stressors including pathogens, pesticides, and environmental pressures (1). These threats have drawn increasing attention to the role of gut microbial communities in maintaining bee health (2–4). The adult honey bee gut hosts a relatively simple, host-adapted microbiota dominated by five core taxa: *Bifidobacterium*, *Lactobacillus* (formerly Firm-5), *Bombilactobacillus* (formerly Firm-4), *Gilliamella*, and *Snodgrassella*, which are primarily acquired through food intake and social interactions within the hive (4, 5). This microbiota supports digestion, detoxification, immunity, and behavioral regulation (4, 6–8), and disturbances to its composition, such as those induced by antibiotics or environmental contaminants, can negatively affect host physiology (9–12). In response, strategies based on probiotic supplementation (13–15), gut homogenate transfer (16), and genome-engineered symbionts (17, 18) have been proposed.

Among the strategies proposed to improve honey bee gut microbiota, supplementation with *Lactobacillus* and *Bifidobacterium* species is considerably promising due to their presence and abundance in the gut and their versatile roles. These taxa, predominantly colonizing the rectum, contribute to the breakdown of dietary polysaccharides and phytochemicals, support nutrient absorption, and produce organic acids such as lactate, succinate, and acetate that reinforce gut homeostasis (19–22). In addition, *Lactobacillus* strains display antagonistic effects against pathogens including *Hafnia alvei* and trypanosomatid parasites (23, 24), and their succinate production has been linked to protection from metabolic dysfunctions in bees (25).

Beyond gut metabolism, *Lactobacillus* and *Bifidobacterium* species have also been implicated in neurophysiological regulation (4, 26). For instance, *Lactobacillus apis* enhances learning and memory by modulating tryptophan metabolism (27), while *Bifidobacterium* and *Bombilactobacillus* species have been associated with increased levels of γ-aminobutyric acid (GABA), a major inhibitory neurotransmitter, through the upregulation of the glutamate receptor gene *gluR-B* (28, 29). Other neuroactive metabolites such as serine and ornithine are also influenced by gut microbes and have been linked to synaptic function and brain energy metabolism (30).

Despite the growing evidence of highlighting the importance of gut symbionts, research investigating the efficacy of probiotics application in honey bees remains inconsistent. Some studies have shown that lactic acid bacterial supplementation can protect against *Paenibacillus larvae* and *Varroa destructor* (14, 31) or enhance immune response following antibiotic treatment (13). However, other studies have found no significant benefits from *Lactobacillus*- and *Bifidobacterium*-based supplementation against *P. larvae* (32, 33). Variations in hive conditions, study design, and delivery methods likely contribute to these contradictory outcomes (31, 34). Engineered gut strains have shown promise in laboratory settings, such as suppressing viral infections and increasing mite mortality (18), but their application under field conditions remains untested. Although various approaches have been proposed to support bee gut health, most studies have conducted experiments exclusively under either laboratory or field conditions, with only a few including both (22, 35–37). However, how bacterial supplementation influences gut physiology through gradual, context-dependent changes across multiple biological levels, such as the microbiome, proteome, and metabolome are not fully understood. This knowledge gap has limited the research field’s ability to comprehensively assess the outcomes of probiotic interventions under different environmental conditions.

To address these gaps, we hypothesized that supplementation with *Lactobacillus* and *Bifidobacterium* strains isolated from the honey bee gut would enhance core symbiont colonization and alter host metabolism and gut protein abundance. We aimed to characterize the consistency and context-specificity of these effects across different environmental conditions by comparing controlled, semi-controlled, and field conditions, in order to identify both shared and environment-dependent patterns in host responses to bacterial supplementation.

## 2. Materials and methods

### Experimental Design

For the cage experiment conducted on 6 August 2022, newly emerged worker bees (*Apis mellifera*) were sourced from a colony in Kývalka, the Czech Republic (49.1913056N, 16.4495556E) and randomly assigned to four groups for the controlled and semi-controlled experimental conditions: control (C) and bacterial supplementation (B) under controlled microbiome conditions to prevent natural microbial acquisition, and control with nestmates (CN) and bacterial supplementation with nestmates (BN) under semi-controlled conditions allowing limited social interaction (Fig.1). Five to ten nestmates from the same colony were introduced, color-marked, and excluded from the final sampling. Each group was housed in plastic cup cages and maintained in an incubator at 34 ± 1 °C in darkness for 10 days (38). All groups received *ad libitum* access to a 50% of sucrose solution, either with or without the bacterial supplement, along with gamma-irradiated pollen. Supplemented groups received a single dose of 1 × 10^9^ CFU per 100 bees of the bacterial supplement on day 1. Once consumed, bees have received only sucrose solutions without further supplementation.

**Figure 1.**
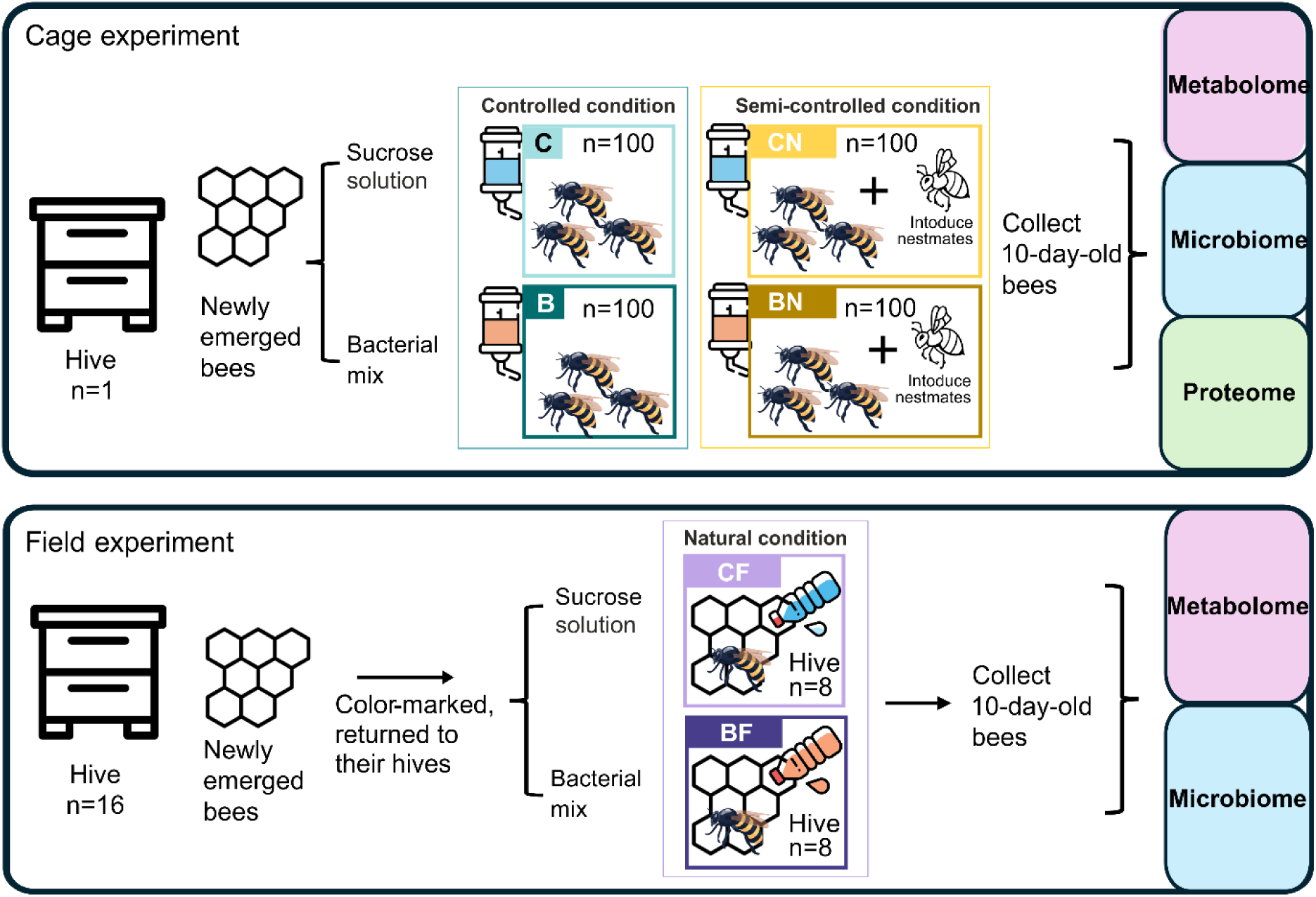
Experimental design and overview of the study. This study comprised two main experimental approaches: cage and field experiments. The cage experiment was conducted under controlled condition (C vs. B) to assess the effects of bacterial supplementation, and semi-controlled condition (CN vs. BN), where nestmates from the same hive were introduced to monitor the influence of social interactions on gut microbiota development together with the bacterial supplementation. The field experiment was carried out in a natural, uncontrolled environment (CF vs. BF) to evaluate these effects under realistic hive conditions.

For the field experiment conducted on 2 August 2022, newly emerged bees (*Apis mellifera*) were collected directly from brood combs at the Postřižín apiary in the Czech Republic (50.2267222N, 14.3778611E), color-marked, and returned to their original hives (Fig. 1). Control hives (CF) received 100 mL of 50% sucrose solution, while bacterial supplemented hives (BF) received 100 mL of 50% sucrose solution containing 4.6 × 10^10^ CFU of the bacterial supplement per hive. Solutions were administered once by trickling between the brood frames. After 10 days, color-marked bees were recollected (335 bees from CF and 326 bees from BF), immediately frozen on dry ice, and stored at −80 °C for further analysis. No visible *Varroa* mite attachment was observed.

### Bacterial supplement preparation

The bacterial supplement consisted of five bee gut-derived strains, including two of *Lactobacillus helsingborgensis*, *Lactobacillus apis*, *Bifidobacterium polysaccharolyticum*, and *Bifidobacterium choladohabitans*, isolated from *Apis mellifera* workers in the Czech Republic. These strains were selected based on a preliminary screening study assessing their antimicrobial properties (39), cultured anaerobically in modified BHI medium, lyophilized, and the bacterial supplement was stored at −80 °C until use. Detailed strain information and culturing protocols are provided in Supplementary methods, section 1.

### Sample dissection and weighing

Both cage and field experiment honey bee gut sample dissections were conducted as previously described (40). Briefly: the venom sacs were removed using sterile tweezers and the gastrointestinal tracts, excluding the crop, were collected. For metabolomics analysis, the wet weight of each gut sample was measured for normalization. For DNA extraction, each bee was individually surface-sterilized following previously described methods (10).

Individual honey bee bodies (excluding the head and venom sac) and the guts (excluding the crop) were placed in separate pre-weighed 2 mL microcentrifuge tubes (VWR, USA), and reweighed. Each measurement was done in triplicate and averaged (Table S1).

### Bacteriome profiling

DNA was extracted from dissected honey bee gut samples using the DNeasy PowerSoil DNA Guts from single bees were processed individually for the cage experiment, while pooled guts from five bees were used for the field experiment. DNA quantity was determined by real-time PCR using Qiagen HotStarTaq DNA Polymerase and primers specific for the V4 region of the 16S rRNA gene, along with a fluorescent probe. Extracted DNA samples were stored at −20°C until further analysis.

Amplicon sequencing was conducted on the V3-V4 region using modified 341-F and 806-R primers with heterogeneity spacers to enhance signal diversity (O. Cinek *et al*., submitted for publication). Each sample was amplified in duplicate, incorporating different adaptor orientations. PCR products were indexed with Nextera primers, purified, normalized, and sequenced using the Illumina MiSeq platform (2 × 250 bp, V2 kit) with 10% PhiX spike-in. The run yielded 14.5 million clusters with >94% Q30 for forward and >86% for reverse reads.

Raw reads were demultiplexed and processed using a custom pipeline: heterogeneity spacers were trimmed, reads merged with USEARCH, and sequence variants inferred using DADA2. Taxonomic classification was performed using the BEExact database (v2023.01.30), and a phylogenetic tree was constructed with FastTree2. Duplicates were assessed via Bray-Curtis ordination and merged. Taxa present in fewer than 80% of samples were removed, and read counts were rarefied to 14,499 per sample. The resulting ASV table, including taxonomic annotations and sample-wise read abundances, is provided in Supplementary Table S2. Detailed sequencing and bioinformatic protocols are provided in the Supplementary Methods, section 2.

### Proteome analysis

Proteomic analysis was performed on gut samples from the cage experiment, following the protocol adapted from Cullen *et al*. (41). Three bees randomly selected from each group. Guts were homogenized in lysis buffer, centrifuged, and protein concentrations were measured using the Qubit™ Protein Assay Kit. Extracted proteins were purified with a 2-D Clean-Up Kit and digested with trypsin in the presence of ProteaseMax™, following reduction and alkylation steps. Peptides were purified using C18 spin columns, vacuum-dried, and resuspended in 2% acetonitrile/0.05% TFA before LC-MS/MS analysis.

Peptides (1 µg per sample) were analyzed using a Q Exactive mass spectrometer coupled to a Dionex Ultimate 3000 nano LC system. Peptide separation was achieved on a C18 column using a 120-minute gradient. The instrument operated in data-dependent acquisition mode, targeting the 15 most intense ions per MS scan (m/z 300–2000) for MS/MS.

Raw data were processed in MaxQuant (v2.4.2.0) using the Andromeda search engine and LFQ normalization, with identification against the *A. mellifera* NCBI protein database (downloaded June 2023). Processed data were filtered and statistically analyzed using Perseus (v2.0.10.0). After imputation of missing values, 977 proteins were retained for further analysis (Table S3). The MS proteomics dataset is available via PRIDE with the identifier PXD066488. Detailed sample preparation, digestion, LC-MS/MS, and data processing protocols are provided in the Supplementary Methods, section 3.

### Metabolome analysis

40 samples from the cage experiment (10 per group: C, CN, B, BN) and 44 from the field experiment (20 CF, 24 BF) were analyzed. For the sample processing, ¹H NMR analysis, and data processing were performed as previously described in Lee *et al* (40). Briefly: Individual bee guts were homogenized with 5 mm zirconium oxide beads in 1 mL methanol using a Retsch MM200 homogenizer (25 Hz, 3 min) (Retsch, DE), ultrasonicated (5 min), and centrifuged (14,000 × g, 10 min, 4 °C). The supernatant was dried using a centrifugal vacuum concentrator at 40 °C (MiVac Duo, UK). This extraction was repeated, and pooled supernatants were dried again.

Dried extracts were reconstituted in 600 µL D₂O, vortexed, and centrifuged (14,000 × g, 5 min). 540 µL of supernatant was mixed with 60 µL of phosphate buffer (1.5 M K₂HPO₄/NaH₂PO₄, pH 7.4, with 5 mM 3-(trimethylsilyl)-2,2,3,3-tetradeuteropropionic acid (TSP) and 0.2% NaN₃ in D₂O). Prepared samples were transferred into 5 mm NMR tubes (Wilmad-LabGlass, USA) in random order for spectral acquisition.

¹H NMR analysis and data processing were performed on a Bruker Avance III spectrometer equipped with a broad band fluorine observation SmartProbe™ with z-axis gradients (Bruker, USA), operating at the ^1^H NMR frequency of 500.18 MHz, as previously described in Lee *et al* (40). Spectra were manually phased in Topspin v3.6.5 (Bruker, USA), with quality assessed based on TSP symmetry and line width (<1 Hz).

Spectral processing was conducted in MATLAB® R2022a using in-house scripts for multipoint baseline correction and binning (δ_H_ 0.5-9.5 ppm), excluding residual water (δ_H_ 4.75-4.90) and methanol (δ_H_ 3.34-3.75). Bin intervals were defined based on spectral annotation using Chenomx NMR Suite v9.0 (Chenomx Inc., Canada), internal libraries. Final intensities were normalized to TSP and sample weight (mg of gut tissue). Metabolite assignments and bin definitions are listed (Table S4).

### Statistics

Statistical analysis of honey bee metabolomics and microbiome data were conducted using R (v4.4.2) (42). The dplyr (v1.1.4) (43) was used as a sub setting tool. Metabolite and taxa comparisons between groups were analyzed using the Wilcoxon rank-sum test, with the Benjamini-Hochberg procedure applied using rstatix (v0.7.2) (44). Principal component analysis (PCA) biplots were visualized with factoextra (v1.0.7) (45). Data visualization was done with ggplot2 (v3.5.1) (46). Results with a *p*-value < 0.05 were considered statistically significant.

Proteomics analysis was performed using Perseus (v2.0.10.0). Normalized intensity values were used for PCA. To identify statistically significant differentially abundant (SSDA) proteins, a two-sample t-test was applied with a significance threshold of *p* < 0.05. Volcano plots were generated by plotting, highlighting proteins that passed both thresholds (log₂ fold change > 1.0; –log₁₀ *p* > 1.3). Hierarchical clustering of SSDA proteins was performed using z-score normalized intensity values, with clustering based on Euclidean distance. Gene Ontology (GO) enrichment analysis was performed using ShinyGo (v0.81) (47) with the *Apis mellifera* dataset based on the STRING protein. Enriched GO terms and pathways (FDR ≤ 0.05) were identified based on SSDA proteins.

## Results and discussion

### *Bifidobacterium* and *Lactobacillus* consistently enriched in supplemented bees

We assessed the impact of bacterial supplementation on honey bee gut microbiota by performing 16S rRNA profiling of all three experiments. In the controlled condition, group C consisted of bees that developed their gut microbiota in a setting designed to restrict external microbial exposure, was dominated by *Bombella* (*q* = 0.018, Table S5), and exhibited elevated levels of other bacterial taxa belonging to non-core microbiota (Figs. 2A and 2B). While group B showed high abundance levels of *Bifidobacterium* and *Lactobacillus* (both *q* = 0.018; Table S5), indicating successful colonization by the supplemented strains (Fig. 2B). This is consistent with previous reports showing that *Lactobacillus* and *Bifidobacterium* can colonize the gut environment of newly emerged or germ-free bees (4, 22).

**Figure 2.**
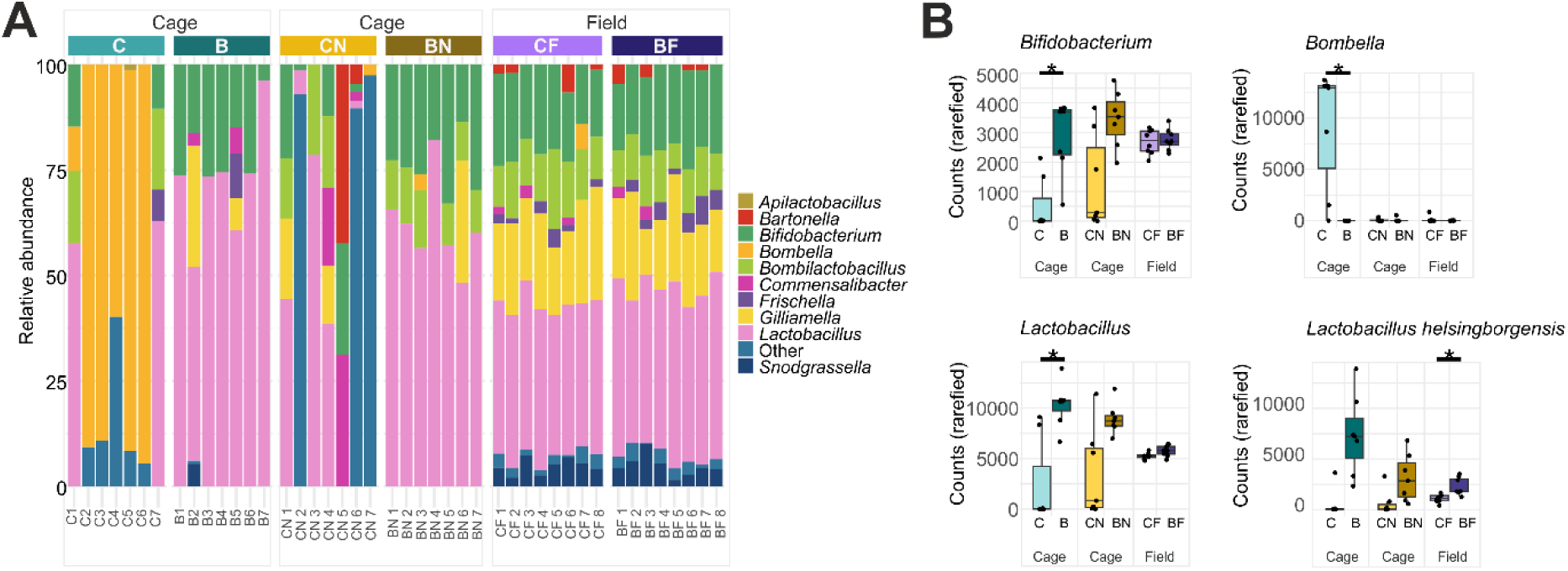
Gut microbiota composition and diversity across experimental groups. **(A)** Relative abundance of bacterial taxa at the genus level in the cage experiment (C, B, CN, BN; *n* = 7 per group) and field experiment (CF, BF; *n* = 8 per group). **(B)** Box plot analysis of statistically significant bacterial genera and species showing differences between control and treatment in at least one experimental condition. Statistical comparisons were performed using the Wilcoxon test, followed by Benjamini-Hochberg procedure (C vs. B, CN vs. BN; *n* = 7 per group; CF vs. BF; *n* = 8 per group). Adjusted *P*-values are indicated with an asterisk (*q* < 0.05). Group abbreviations: C, control (cage); B, bacterial supplementation (cage); CN, control with nestmates (cage); BN, bacterial supplementation with nestmates (cage); CF, control (field); BF, bacterial supplementation (field).

In the semi-controlled experiment, BN supplementation promoted a more consistent establishment of *Lactobacillus* and *Bifidobacterium* compared to CN. However, this difference was not statistically significant (Fig 2B, Table S4), likely due to high inter-individual variability (Fig 2A), which may reflect inconsistent microbial transfer from older nestmates (28, 48). It is also important to note that in the cage experiments, honey bees do not defecate (49, 50), resulting in minimal microbial turnover compared to hive conditions. Consequently, supplemented strains are more likely to be retained within the digestive tract.

In the field experiment, the bacteria supplemented group (BF) received the same supplement as in the cage trials. Although genus-level profiles were broadly similar between BF and CF (Fig. 2A), species-level analysis revealed a significant enrichment of *Lactobacillus helsingborgensis* in BF (*q* = 0.035; Fig 2B, Table S5). Other administered strains did not show significant differences. This selective colonization suggests that *L. helsingborgensis*, which comprised approximately one-third of the living bacterial cells of the supplement, was uniquely capable of establishing in the competitive and dynamic environment of the field. Supplementation induced marginal changes in the relative abundance of several taxa, including *Lactobacillus* spp., *Bifidobacterium* spp., *Gilliamella* spp., and *Commensalibacter* with some increasing and others decreasing; however, none of these effects remained statistically significant after Benjamini-Hochberg procedure (Table S5).

The dominance of *L. helsingborgensis* in BF underscores its ecological resilience and compatibility with the native gut microbiota. This species is a core member of the honey bee gut microbiota, known for its role in carbohydrate fermentation (51), and has demonstrated stress resilience under diverse chemical exposures (52, 53). More broadly, *Lactobacillus* species contribute to immune regulation in the honey bee (4), with *L. helsingborgensis* and *L. apis* reported to reduce larval mummification (54), and inhibit *Paenibacillus larvae* and *Melissococcus plutonius* growth *in vitro* (39, 55, 56). However, such effects may not consistently translate to field conditions (32, 33), where interactions with native microbiota, environmental filtering, and niche competition can alter outcomes.

Overall, these results suggest that while laboratory supplementation leads to reliable colonization by core taxa, real-world applications require consideration of ecological compatibility. Strains like *L. helsingborgensis* that demonstrate both functional benefits and field persistence may serve as promising candidates for probiotic development in honey bees.

### Bacterial supplementation influence on host gut proteome responses

To investigate the effects of bacterial supplementation on the honey bee gut tissue proteome, we compared protein expression of the cage experiments using label-free quantification. PCA revealed clear group separation in both comparisons, indicating that supplementation elicited distinct shifts in protein abundances (Fig. 3A top). The BN group exhibited greater inter-individual variability along the PC1 axis, suggesting a more heterogeneous response compared to the relatively consistent clustering seen in B (Fig. 3A bottom).

**Figure 3.**
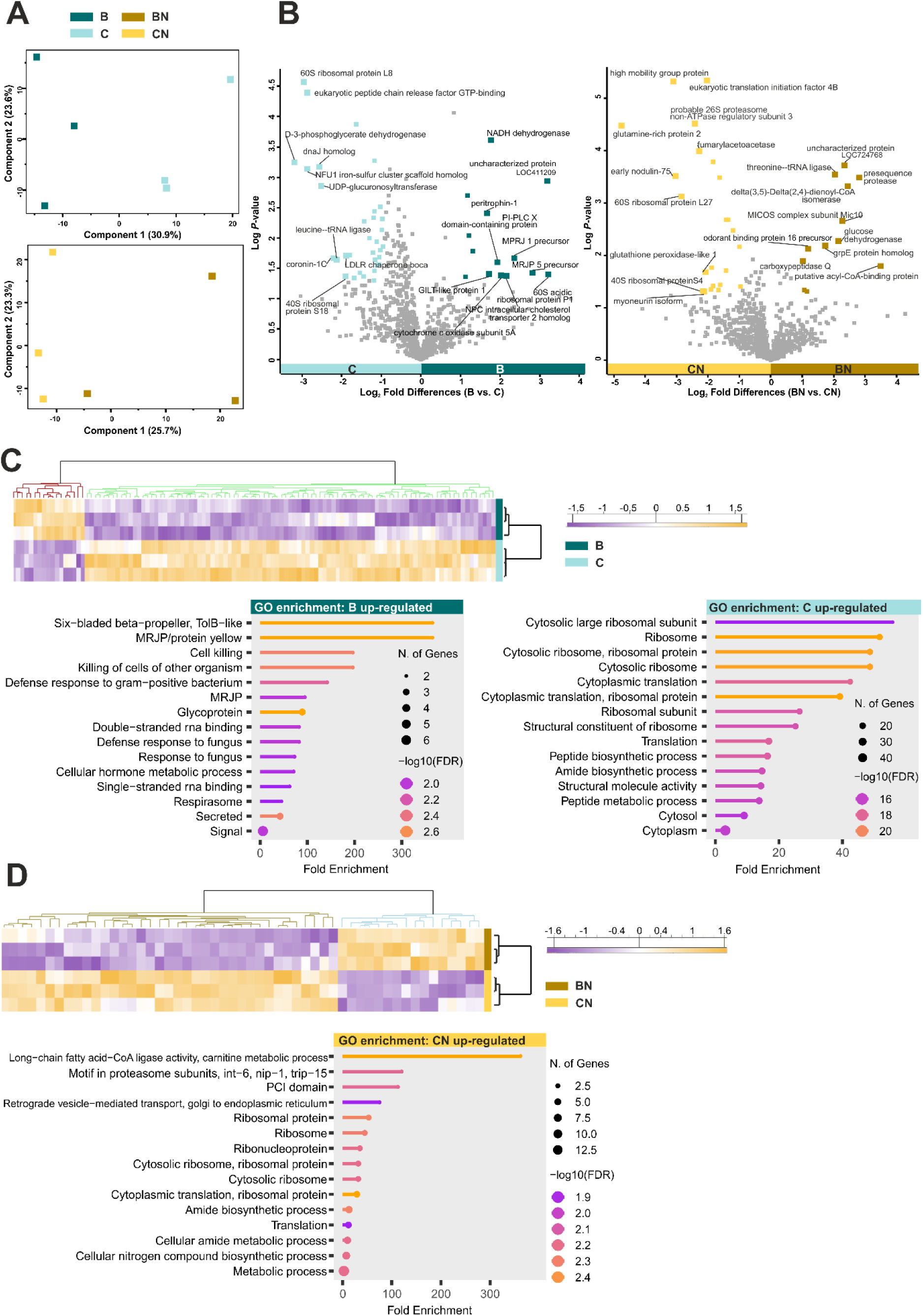
Label-free quantitative mass spectrometry analysis of host gut proteome differences in honey bees following bacterial supplementation and social interaction conditions **(A)** Principal component analysis (PCA) of honey bee host gut proteomes comparing B vs. C (top) and BN vs. CN (bottom). **(B)** Volcano plots illustrate differential protein abundances in comparisons between B vs. C (left) and BN vs. CN (right). Differential abundance was determined using a threshold of –log₁₀(p) ≥ 1.3 and |log₂ fold change| ≥ 1.0. Significantly more and less abundant proteins (n = 47 in B vs. C; n = 35 in BN vs. CN) are highlighted, with the top ten proteins labelled based on fold-change in each comparison. Non-significant proteins are shown in grey. **(C)** Hierarchical clustering heatmap of statistically significant differentially abundant (SSDA) proteins in B vs. C, alongside with GO enrichment analysis of upregulated proteins in each group, showing the top 15 significantly enriched pathways. **(D)** Hierarchical clustering heatmap of SSDA proteins in BN vs. CN, with GO enrichment analysis of upregulated proteins in the CN group. No significant GO enrichment was detected in the BN group. Group abbreviations: C, control (cage); B, bacterial supplementation (cage); CN, control with nestmates (cage); BN, bacterial supplementation with nestmates (cage).

In the B vs. C comparison, 113 SSDA proteins were identified, 47 surpassing both fold-change (FC) and significance thresholds (Fig. 3B left, Table S6). These proteins also showed clear group-specific clustering in hierarchical heatmaps (Fig. 3C). The B group displayed increased abundance of major royal jelly protein precursors MRJP1 and MRJP5 (FC: +5.1 and +7.1, respectively) compared to C, which contributed to the enrichment of GO categories such as MRJP, defense response, and cell killing (Fig. 3C, Table S7). These results align with prior findings showing that mono-colonization by *Lactobacillus* and *Bombilactobacillus* induces MRJP expression in the honey bee brain and modulates neurotransmitter levels (27, 29), supporting the concept of a microbiota-gut-brain axis. MRJP1 has been linked to nutrient processing, caste development, and antimicrobial functions (57, 58), while MRJP5 is primarily known for its defensive role (59). Interestingly, MRJP expression is typically reduced in *Varroa*-infested and *Nosema*-infected bees, where energetic stress impairs protein secretion (38)(60). Overall, these patterns suggest that bacterial colonization supports early immune priming and nutritional signaling.

Group B showed a decreased abundance of ribosomal and translation-associated proteins compared to the group C, including 60S ribosomal protein L8 (FC: −7.8) and 40S ribosomal protein S18 (FC: −3.7) (Table S6). These proteins, enriched in the C group, were associated with GO terms related to cytoplasmic translation and peptide biosynthesis (Fig. 3C, Table S7) and are essential not only for translation but also for ribosome assembly, structural integrity, and stress response (61–63). A similar transcriptional response was reported in bees exposed to sublethal imidacloprid (64), suggesting that microbial deprivation may lead to comparable compensatory mechanisms in the gut. Ribosomal genes have also been shown as stable reference markers during viral and dsRNA challenges (65). Given their consistent expression in the hypopharyngeal glands of nurse bees (66), their decreased abundance in B may reflect reduced translational demand or tissue remodeling in response to symbiont absence.

In the BN vs. CN comparison, 54 SSDA proteins were detected, with 35 meeting fold-change and significance thresholds (Fig. 3B right, Table S6). Hierarchical clustering confirmed distinct group-wise proteomic profiles (Fig. 3D). No GO pathways were significantly enriched in the BN group; however, manual inspection revealed functionally coherent protein changes. These proteins with increased abundance were associated with lipid metabolism (acyl-CoA-binding protein, FC: +11.2; delta(3,5)-delta(2,4)-dienoyl-CoA isomerase, FC: +5.5) (67), as well as mitochondrial proteostasis and structure (GrpE homolog FC: +3.3, Mic10; FC: +4.9) (68, 69) (Table S6). These findings suggest that bacterial supplementation combined with social microbiota input may promote mitochondrial adaptations aligned with gut metabolic needs. Still, the absence of GO-level enrichment limits interpretation, making it unclear whether these shifts reflect a coordinated biological program or independent responses to colonization.

Conversely, BN bees showed lower abundance of several ribosomal and stress-associated proteins, including 60S ribosomal protein L27a (FC: −7.2), 40S ribosomal protein S4 (FC: −4.5), heat shock protein cognate 3 precursor (FC: −1.3), and UBX domain-containing protein 4 (FC: −3.6), all of which were more abundant in the CN group (Table S6). These proteins contribute GO categories related to cytoplasmic translation and peptide biosynthesis, and endoplasmic reticulum (ER)-associated and response to unfolded protein pathways (Table S7).

These pathways are central to proteostasis and cellular protein quality control, often triggered by toxin exposure, misfolded protein accumulation, or metabolic imbalance (70, 71). A previous study has shown that honey bee guts mount a strong unfolded protein response under ER stress (72), and our findings suggest that bacterial supplementation with nestmates may reduce such stress, as reflected by the lower abundance of these markers in BN bees.

Taken together, the data indicate that bacterial supplementation results in group-specific proteomic changes in the honey bee gut. The B group displayed an MRJP-enriched signature, while the BN group showed a distinct profile characterized by mitochondrial and lipid-related adaptations. Both B and BN groups showed decreased levels of translation-related proteins, and the BN group also exhibited a reduced abundance of ER stress-associated proteins, which were more prominent in the C and CN bees.

### Gut metabolic alterations under bacterial supplementation and consistent trends across experimental conditions

To evaluate metabolic alterations induced by bacterial supplementation, we conducted ^1^H NMR-based metabolomics in all three experiments. A total of 57 metabolites were identified, including amino acids, amines, carbohydrates, lipids, purine/pyrimidine metabolism, sulfur compounds, phenolic acids, and carboxylic/hydroxy/fatty acids (Table S4). The PCA biplots of the top 20 contributing metabolites revealed clear separation between bacteria supplemented and control groups in both cage comparisons (Figs. 4A and 4B). Although intra-group variability was higher in the field comparison, some separation was evident (Fig. 4C). Metabolites such as *N*-acetylglucosamine (GlcNAc), acetate, leucine, and isoleucine consistently contributed to group discrimination across all conditions (Figs. 4A-C).

**Figure 4.**
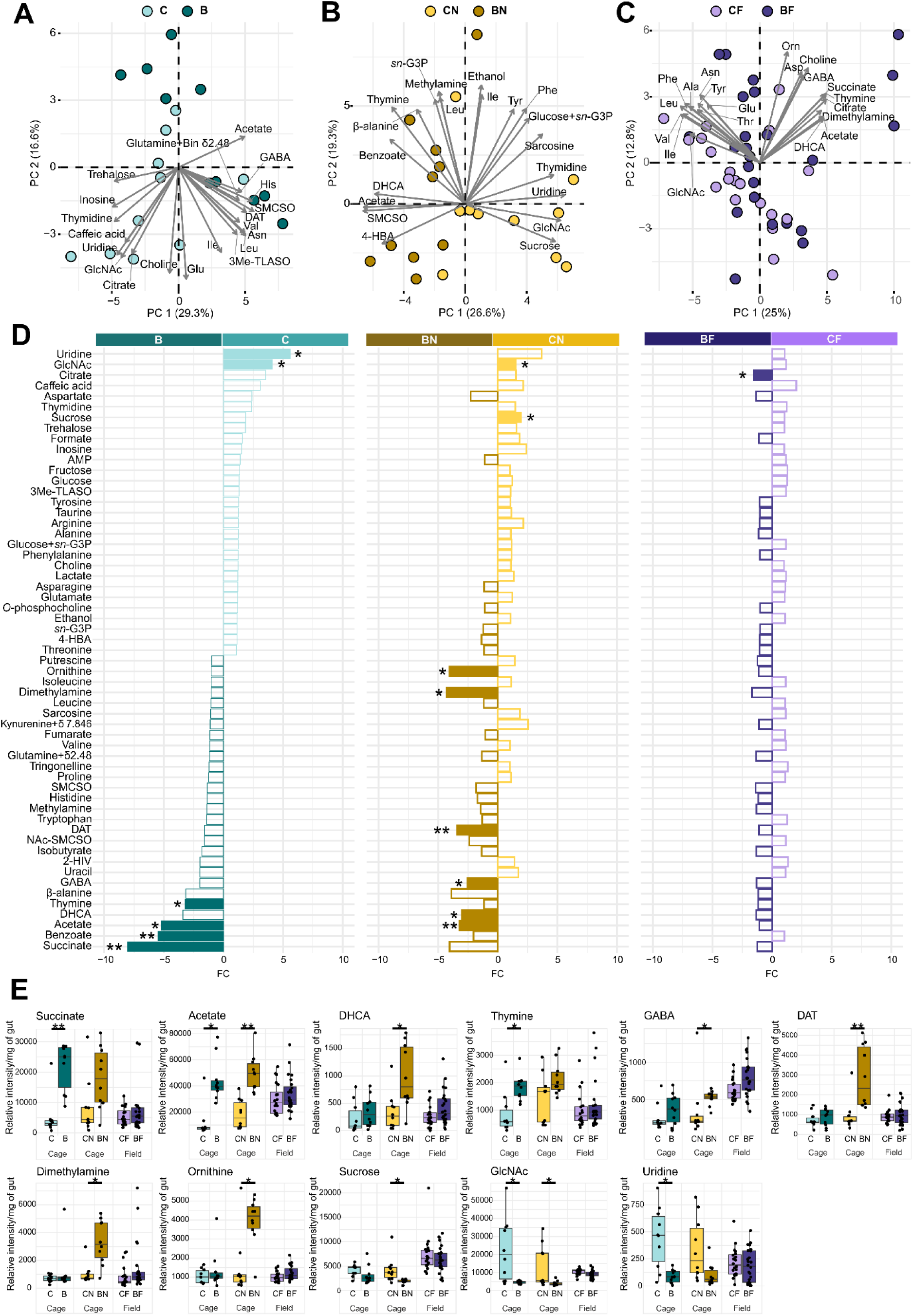
Metabolic profiling of honey bee gut samples across experimental groups in cage and field experiments **(A–C)** Principal component analysis (PCA) biplots showing metabolomic composition in honey bee gut samples for C vs. B (cage), CN vs. BN (cage), and CF vs. BF (field). Gray arrows represent the top 20 contributing variables based on loading scores. **(D)** Fold-change analysis of 57 metabolites from honey bee gut, presented as median fold changes of groups. Statistically significant adjusted *P*-values are indicated with an asterisk mark and a bar with filled colors (* *q* < 0.05, ** *q* < 0.01, no mark = ns). **(E)** Box plot analysis of statistically significant metabolites between C vs. B, CN vs. BN, and CF vs. BF. A total of 11 metabolites were selected based on their consistent trends across both cage and field experiments. Statistically significant adjusted *P*-values are shown in asterisks mark (* *q* < 0.05, ** *q* < 0.01, no mark = ns). Statistical comparisons were performed using the Wilcoxon test, followed by Benjamini-Hochberg (BH) procedure (C, B, CN, BN; *n* = 10 per group; CF, *n* = 20; BF, *n* = 24). Group abbreviations: C, control (cage); B, bacterial supplementation (cage); CN, control with nestmates (cage); BN, bacterial supplementation with nestmates (cage); CF, control (field); BF, bacterial supplementation (field).

To compare gut metabolic responses across the three experimental conditions, we ranked metabolites by fold change in the controlled experiment and compared trends across semi-controlled and field experiments. The two cage experiments showed similar patterns, with 19 metabolites elevated in treatments and 22 in controls (Fig. 4D). However, the field setting shared limited overlaps, with citrate being the only significantly altered metabolite (*q* = 0.048, Fig. S1), and its direction of change differed from cage results (Fig. 4D, Table S8). This contrasts with its depletion in colonized bees under laboratory conditions (22, 73), suggesting that the divergence is likely due to environmental filtering or different experimental settings. Further analysis revealed 11 metabolites showing consistent trends across the three experiments that are significantly different in at least one experiment (Fig. 4E). These patterns suggest that subtle but biologically meaningful metabolic shifts may also occur under hive conditions, although their detectability is likely obscured by environmental noise and natural biological variation in the field.

### Bacterial supplementation and microbial fermentation products

Succinate and acetate, key fermentation metabolites, were significantly elevated in all treatment groups (Fig. 4E). Succinate increased by 8.1-fold in B vs. C (*q* = 0.004) and 4.1-fold in BN vs. CN (marginally significant; *q* = 0.073), while one of short chain fatty acids (SCFAs), acetate increased 5.2-fold (B), 3.2-fold (BN), and 1.1-fold (BF) over respective controls (*q* = 0.014, *q* = 0.002, and ns; Tables S8-S9). These metabolites typically produced by *Lactobacillus* and *Bifidobacterium* (19, 21, 74), likely reflect active fermentation and microbial integration. Their accumulation is known to support anaerobiosis and microbiota stability (4, 22), and also serve as substrates for cross-feeding microbes (75, 76). In addition to microbial functions, succinate has been implicated in regulating host metabolism through gut gluconeogenesis and insulin-like signaling (25). These metabolites may serve not only as fermentation byproducts but also as signaling molecules influencing host physiology. The increased abundance of glucose dehydrogenase (FC: + 2.5 in B; Table S6) suggests increase host carbohydrate oxidation, potentially facilitating microbial access to fermentable sugars or complementing microbial carbohydrate metabolism (77). Concurrently, increased abundance of NADH dehydrogenase (FC: +3.4 in B; Table S6), a key enzyme in the mitochondrial respiratory chain indicates enhanced oxidative phosphorylation (78), possibly driven by increased availability of microbial-derived fermentation metabolites. Together, these changes suggest coordinated microbial-host metabolic interactions, where bacterial fermentation reshapes host energy and sugar metabolism.

We observed a marginally significant increase in body weight in the B group compared to C (*p* = 0.075; Fig. S2, Table S10), consistent with a report linking microbial colonization to weight gain via bacterial metabolism and SCFAs production (16). However, this pattern was absent in the BF and in BN bees, indicating that host weight responses are context-dependent (79) (Fig. S2, Table S10).

BN showed significantly elevated levels of other microbial fermentation products, including dihydrocaffeic acid (DHCA; 3.0-fold, *q* = 0.027), dimethylamine (4.3-fold, *q* = 0.010), and desaminotyrosine (DAT; 3.4-fold, *q* = 0.009) (Fig. 4E, Tables S8-S9). Lactic acid bacteria can convert caffeic acid to DHCA, aiding to metabolism of plant phytochemicals. Similar changes have been documented in microbiota-colonized bees (22, 80, 81). Dimethylamine is likely derived from nitrogenous precursor metabolism, as seen in *Enterobacteriaceae* and *Lactobacillus* (82–84). DAT, a microbial catabolite of tyrosine and flavonoids, is recognized for its immunomodulatory effects in mammals (85–87), though it remains undescribed in honey bee gut metabolites studies (16, 22). These increased metabolites likely reflect microbial transmission from nestmates and synergistic degradation alongside supplemented strains. Although the B group showed similar trends, the effects were not statistically significant (Table S8), highlighting the role of social interaction in stabilizing or amplifying microbial function.

### Increased neuroactive metabolites in bacteria-supplemented bees

Beyond fermentation, bacterial supplementation was associated with increases in neuroactive metabolites, including GABA, β-alanine, and ornithine (Fig. 4E, Fig. S3). Especially in the BN vs. CN comparison, GABA and ornithine were significantly higher in BN than in CN (GABA: 2.5-fold, *q* = 0.027; ornithine: 4.0-fold, *q* = 0.049; Fig 4E, Tables S8-9). These compounds have been implicated in neurotransmission, behavior, and sensory modulation in honey bees (26, 29, 30).

Notably, bacterial supplementation led to an increase in GABA levels, a key inhibitory neurotransmitter involved in sensory processing, learning, and motor control in honey bees, (26, 88, 89). Behavioral studies show that GABA administration enhances learning and memory when paired with a reward, but impairs learning acquisition when ingested prior to training (88). Additionally, GABA injection influenced grooming behavior, and disruption of GABA signaling impaired motor coordination (89). Previous studies have similarly reported elevated GABA levels and modulation of brain gene expression following supplementation with *Bifidobacterium* and *Bombilactobacillus*, suggesting a potential gut–brain axis (28, 29). These findings indicate that bacterial supplementation may elicit consistent neurochemical responses. Although β-alanine narrowly missed statistical significance after multiple testing correction (*q* = 0.056; Table S8), the trend was reproducible across all experiments (Fig. S3). As a GABA receptor agonist found in the bee brain, β-alanine is associated with sensory learning and social behavior (26, 30, 88). We did not observe significant changes in tryptophan (Fig S3, Table S8), which is known to be converted into behavior-modulating indole compounds by *L. apis* (27). Ornithine, elevated in the same groups, is also implicated in social behavior (26, 30). Supporting these metabolic findings, proteomic analysis revealed elevated abundance of an odorant-binding protein (FC: +2.3 in BN; Table S6), a molecule involved in chemosensory perception (90). Taken together, these elevated neuroactive metabolites suggest that *Lactobacillus* and *Bifidobacterium* supplementation may boost gut-derived neuromodulators, potentially influencing brain signaling and behavior.

### Lower carbohydrate metabolites in bacteria-supplemented bees

The treatment groups (B and BN) showed lower concentrations of uridine, GlcNAc, and sucrose compared to their respective controls (C and CN). Specifically, GlcNAc concentrations were 4.1-fold lower in B compared to C and 1.5-fold lower in BN compared to CN (all *q*-values = 0.049; Fig. 4E, Tables S8 and S9). Sucrose concentrations were 1.9-fold lower in both treatment groups relative to their controls (B vs. C, ns; BN vs. CN, *q* = 0.010; Tables S8 and S9), and uridine showed a similar pattern, with a 5.6-fold decrease in B vs. C (*q* = 0.032) and a 3.7-fold decrease in BN vs. CN (marginally significant; *q* = 0.070) (Fig. 4E, Tables S8 and S9). Sucrose was the major dietary source in this study, and this patterns are likely due to fermentation by supplemented *Lactobacillus* and *Bifidobacterium*, which metabolize dietary sugars into organic acids (19, 22). This is further supported by lower levels of other carbohydrate metabolites in treatment groups, including glucose, fructose, and trehalose (Fig. S3).

Uridine and GlcNAc are intermediates in the hexosamine biosynthetic pathway, which produces UDP-GlcNAc, a key substrate for protein glycosylation, chitin synthesis, and epithelial maintenance (91–93). Their lower concentrations in the B and BN groups compared to C and CN may reflect more efficient microbial recycling, reduced epithelial stress, or stabilized gut metabolism resulting from bacterial supplementation. Elevated GlcNAc levels have been associated with epithelial stress or impaired recycling, as seen during chalkbrood infection, where chitin degradation releases GlcNAc into the bee gut (94). Although no infection was present in this study, similar signatures in controls may reflect dysbiosis or peritrophic matrix remodeling under stress. In the absence of microbial recycling, these metabolites may accumulate, signaling gut dysfunction or delayed epithelial renewal (95). Thus, the observed patterns in treatment groups may indicate balanced host-microbe interactions and reduced accumulation of stress-associated metabolites compared to controls.

Proteomic data support this interpretation, showing reduced abundance of proteins related to translation, peptide biosynthesis, ER-associated degradation, and unfolded protein response in B and BN compared to their controls (Table S7). The reduced abundance of these stress-related proteins in treatment groups aligns with the lower concentrations of GlcNAc and uridine, metabolites involved in glycoprotein biosynthesis and epithelial maintenance. This parallel decrease suggests that bacterial supplementation may stabilize gut epithelial function and reduce the activation of ER stress pathways compared to controls.

### Implications and limitations

This study provides a multilayer view of how bacterial supplementation alters honey bee gut biology across three experimental conditions. Several limitations should be considered. First, the 10-day duration limited assessment of long-term colonization and outcomes relevant to beekeeping. Second, although environmental variability was anticipated, the field experiment introduced uncontrolled factors such as forage diversity and microbial exposures, contributing to higher data variability. Third, proteomic analysis was limited to the cage experiment, restricting insights into protein-level responses under natural conditions. Lastly, functional outcomes like immunity or behavior were not evaluated, which would have strengthened connections between molecular changes and physiological impact. Future studies should assess the long-term effects and practical benefits of this bee-derived supplement under field conditions. Despite these limitations, the study provides a valuable framework for understanding context-dependent host-microbe interactions. These insights can inform probiotic design, improve bee health strategies, and guide future research into the mechanisms linking gut microbiota to host physiology.

## Conclusion

This study demonstrated that bee-derived bacterial supplementation influenced gut microbiota composition, protein abundance, and host metabolism in a context-dependent manner. In cage experiments, *Bifidobacterium* and *Lactobacillus* strains established robustly, while field experiments showed more variable host responses. However, *Lactobacillus helsingborgensis* remained significantly enriched in supplemented bees, indicating ecological resilience and environmental filtering. Despite environmental differences, several features were conserved: *L. helsingborgensis* emerged as a promising probiotic candidate due to its persistence and functional impact. Proteomic analysis revealed increased levels of MRJP precursors and reduced ribosomal and ER stress-related proteins in supplemented bees, suggesting decreased epithelial stress and altered translational demand. Supplemented bees also exhibited higher concentrations of microbial fermentation products and neuroactive metabolites, with lower levels of carbohydrate-associated compounds compared to controls. Together, these findings provide a system-level view of host-microbe interactions and support the development of microbiome-informed strategies to improve honey bee resilience and overall pollinator health.

## Acknowledgement

We thank Jef Engelen for his assistance with sample collection in the field and with the extraction procedures for the NMR analysis. We also gratefully acknowledge the staff of the Bee Research Institute at Dol for their invaluable help with thorax coloring and the collection of honey bee samples. ChatGPT-4o (OpenAI) was employed for language editing. We also thank Dr. Zachary Lavengood for his assistance with language editing. BEI Resources (Manassas, VA) are gratefully acknowledged for providing us with the Microbial Mock Community DNA (HM-276D). Icons used in Figure 1 were obtained from Flaticon.com.

## Funding

This project was funded by the Ministry of Agriculture of the Czech Republic (Project No. QK21010088). This work utilized the NMR facility of the METROFOOD-CZ Research Infrastructure (https://metrofood.cz), supported by the Ministry of Education, Youth and Sports of the Czech Republic (Project No. LM2023064). The Q-Exactive Quantitative Mass Spectrometer at Maynooth University was funded under the Science Foundation Ireland Research Infrastructure Call 2012; Grant Number: 12/RI/2346(3).

## Data Availability

The data underlying this article are available in the supplementary material.

